# Sick plants in grassland communities: a growth-defense trade-off is the main driver of fungal pathogen abundance and impact

**DOI:** 10.1101/806299

**Authors:** Seraina L. Cappelli, Noémie A. Pichon, Anne Kempel, Eric Allan

**Affiliations:** Institute of Plant Sciences, University of Bern, Altenbergrain 21, 3013 Bern, Switzerland

**Keywords:** fungi, infection, pathogen, growth-defense trade-off, grassland, biodiversity experiment, exclusion experiment, specific leaf area, nitrogen, diversity, fungicide

## Abstract

Aboveground fungal pathogens can substantially reduce biomass production in grasslands. However, we lack a mechanistic understanding of the drivers of fungal infection and impact. Using a global change biodiversity experiment we show that the trade-off between plant growth and defense is the main determinant of fungal infection in grasslands. Nitrogen addition only indirectly increased infection via shifting plant communities towards more fast growing species. Plant diversity did not decrease infection, likely because the spillover of generalist pathogens or dominance of susceptible species counteracted dilution effects. There was also evidence that fungal pathogens reduced biomass more strongly in diverse communities. Further, fungicide altered plant-pathogen interactions beyond just removing pathogens, probably by removing certain fungi more efficiently than others. Our results show that fungal pathogens have large effects on plant functional composition and biomass production and highlight the importance of considering changes in pathogen community composition to understand their effects.

## INTRODUCTION

Pathogenic fungi are omnipresent in the environment and have large impacts on their hosts (Fisher *et al*. 2012). Many studies have looked at species-specific (fungus-plant) interactions (e.g. Thrall & Burdon 2003, Roscher *et al*. 2007), however, only a few experiments have investigated fungal pathogens in whole plant communities, by manipulating pathogen access to their hosts (Peters & Shaw 1996; Mitchell 2003; Allan *et al*. 2010; Borer *et al*. 2015; Heckman *et al*. 2017). These studies show fungal pathogens can have large top-down effects, even reducing grassland biomass production as much as insect herbivores (Allan *et al*. 2010; Seabloom *et al*. 2017). However, effects can be context dependent and factors such as plant species composition (Mitchell *et al*. 2002; Rottstock *et al*. 2014) or environmental factors (Mitchell *et al*. 2003) can determine infection rates and pathogen impact. Increasing our knowledge about causes and consequences of fungal pathogens is important to predict effects of global change, e.g. nitrogen enrichment. Nitrogen input can alter pathogen infection (Burdon *et al*. 2006) but the mechanisms by which it does so and the consequences for pathogen abundance and impact are poorly understood.

Key determinants of infection success and consequences of infection are related to pathogen transmission, host resistance and host tolerance to infection (as discussed in detail by Keesing *et al*. (2006)). Transmission and resistance should directly influence the observed levels of infection, while tolerance should alter the negative consequences of infection for fitness or biomass production and, if a tolerant species is a good reservoir host, the infection levels in other species (spillover, Power & Mitchell (2004)). All of these factors can be influenced by environmental variables and might trade-off with each other.

Pathogen resistance and tolerance are linked to plant growth strategy. Plants face a trade-off between growth and enemy defense (***growth-defense trade-off***). Plant species adapted to resource-rich environments grow fast but are often less defended against enemies, including herbivores (Endara & Coley 2011; Lind *et al*. 2013) and fungal pathogens (Blumenthal *et al*. 2009; Liu *et al*. 2017). Fast-growing species are likely to better tolerate enemies, as the loss of plant tissue can easily be replaced (Gianoli & Salgado-Luarte 2017). Hence, plant communities dominated by fast-growing species should display higher pathogen infection but lose less biomass to pathogens than communities dominated by slow-growing plants from resource-poor environments. The leaf economics spectrum distinguishes these strategies and is indicated by several functional traits. Slow-growing species with long-lived, structurally expensive leaves, with low nutrient contents occur at one end of the spectrum and fast-growing species with a high turnover of short-lived, nutrient-rich leaves at the other end (Wright *et al*. 2004). Some of these traits are also directly related to resistance to natural enemies, e.g. leaf nutrient concentrations (Robinson & Hodges 1981). Although the growth-defense trade-off hypothesis is well supported for individual plant-pathogen interactions, we know little about how it scales up to whole plant communities and its importance relative to other drivers of infection.

Another possible driver of infection is the nutrient supply in plant communities. High nitrogen supply can lead to decreased infection resistance with fungal pathogens (***nitrogen-disease hypothesis***, Dordas 2008). Nitrogen disease effects are mainly known from agriculture, while studies in natural ecosystems show more variable results (Mitchell *et al*. 2003; Veresoglou *et al*. 2013). This variation may be partly because nitrogen enrichment can also have complex indirect effects on fungal infection. Nitrogen enrichment often reduces plant species richness and changes plant functional composition by promoting fast-growing over slow-growing species (Bobbink *et al*. 2010; De Schrijver *et al*. 2011; Isbell *et al*. 2013) both of which could indirectly alter fungal infection and its consequences. However, nitrogen could also directly lead to healthier and more tolerant plants. To mechanistically understand nitrogen effects on fungal infection, studies therefore need to assess the direct and indirect effects independently.

Plant diversity can also be a key driver of pathogen infection, through different mechanisms. Pathogen infection has been shown to decrease with greater host diversity in grasslands through changes in plant abundances (e.g. Mitchell *et al*. 2002; Liu *et al*. 2016; Rottstock *et al*. 2014; but see Halliday *et al*. 2017), which reduces host-pathogen transmission (***host dilution hypothesis***) (Civitello *et al*. 2015). However, other studies showed that diverse communities are more infected, potentially due to the spillover of generalist pathogens between plant species or due to an increase of host-density independent pathogens, such as vector-transmitted ones (Power & Mitchell 2004; Halliday *et al*. 2017). The impact of plant diversity on pathogen infection may therefore depend on the relative abundance of specialist and generalist pathogens and on their transmission mode. Plant diversity might also change pathogen community composition by selecting for more generalist species (Thrall *et al*. 2007) and this could potentially alter the impact of pathogens if specialists and generalists differ in their virulence (Leggett *et al*. 2013). However, relatively little is known about the impact of pathogens in low and high diversity plant communities (but see Seabloom *et al*. 2017; Halliday *et al*. (2017)).

Changes in plant functional composition, diversity and nutrients could all affect pathogen communities by changing plant biomass and thereby altering microclimatic conditions. Pathogens generally grow better in warmer and humid conditions, but this varies between pathogen groups (Barrett *et al*. 2009). The availability of free water if often an important driver of infection (Bregaglio *et al*. 2013; Chen *et al*. 2014; Sun *et al*. 2017; Bradley *et al*. 2003), suggesting that the microclimatic humidity is important. Further, increased temperature may promote overall pathogen infection (Liu *et al*. 2016), but again, different groups of fungal pathogens may react differently (Helfer 2014). We therefore lack a good understanding of how temperature and humidity impact different pathogen groups and how these effects relate to other drivers of infection.

The impact of pathogen infection on plant communities mainly depends on the resistance and the tolerance of plants. Impact can be assessed in two ways: comparing plots with and without fungicide, and, assessing the amount of fungal infection in a plant community and relating it to the biomass produced. The first approach would be ideal if fungicide reduced infection to zero. However, most fungicides do not completely wipe out all infection, and might be selective for certain fungal groups (e.g. Parker *et al*. 2015; Karlsson *et al*. 2014), changing fungal community composition. For example, if a fungicide is selective against the rather specialized rusts, then the fungicide might cause a shift from specialized to more generalist fungal communities. The second approach, relating infection and biomass, allows for more quantitative comparisons. However, here the direction of causality is hard to establish, as higher plant biomass might also lead to higher fungal infection, obscuring the relationship. It is therefore advantageous to use both methods, however, no previous studies have done so.

Here we tested the relative importance of nitrogen, plant diversity and functional composition as drivers of fungal pathogen abundance, in an experiment that manipulated these variables factorially (Figure S1, Table S1). Specifically, we tested the growth-defense trade-off hypothesis, the nitrogen-disease hypothesis and the dilution-effect hypothesis (Figure 1). Further, we assessed the fungal impact on plant biomass by comparing biomass from plots with and without fungicide, and by relating plant biomass to infection intensity in the same plots. In addition, we tested if our experimental treatments altered pathogen abundance and impact through changes in microclimatic conditions.

**Figure 1.**
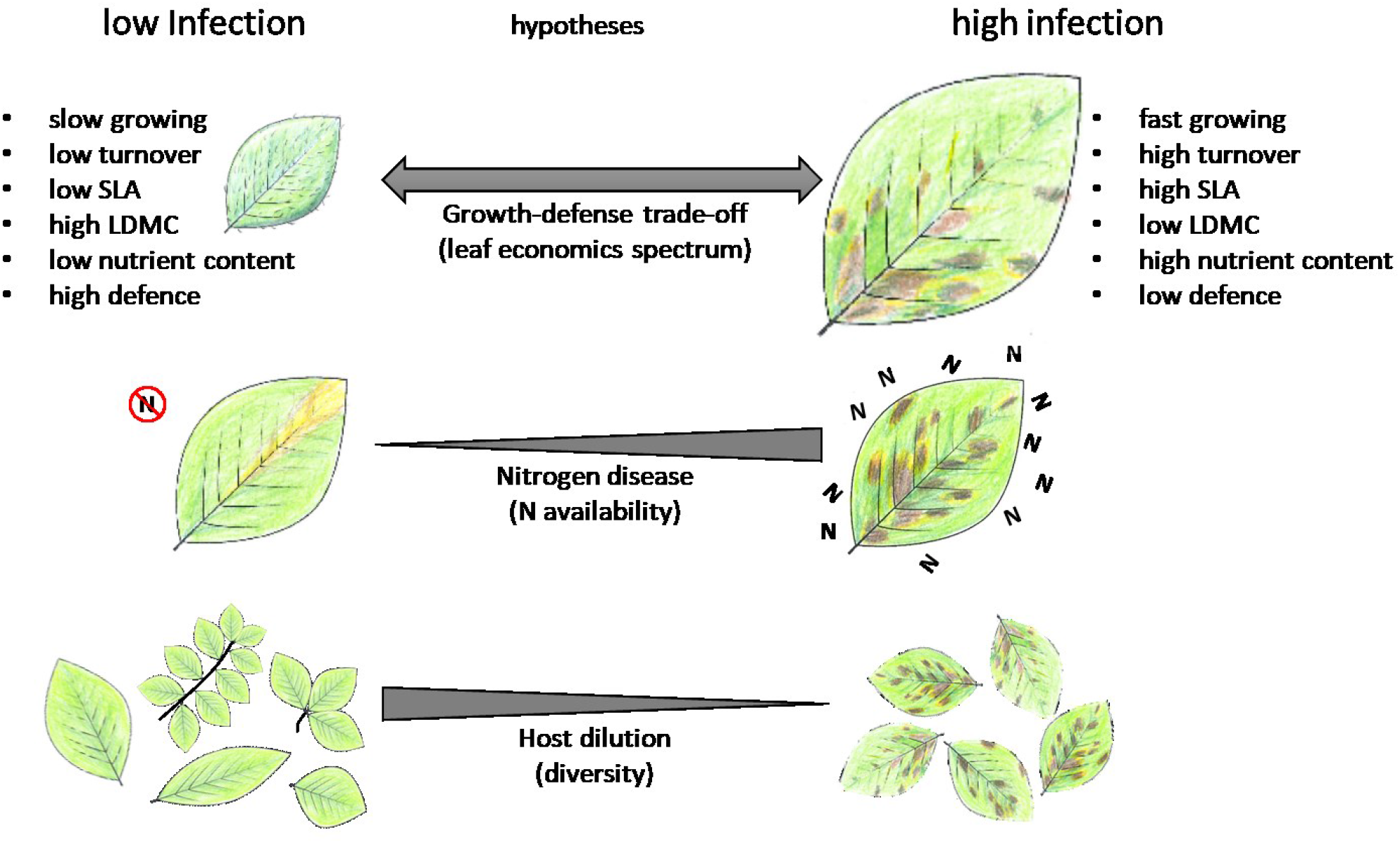
Overview over the main hypotheses which we tested. Growth-defense trade-off hypothesis: Plant species adapted to resource-rich environments and able to compete well under nutrient rich conditions are often less defended against natural enemies (Blumenthal *et al*. 2009; Liu *et al*. 2017). The growth strategy is definded by the leaf economics spectrum (Wright *et al*. 2004), which has been linked to certain disease resistance mechanisms (Cronin *et al*. 2014; Cronin *et al*. 2010; Huot *et al*. 2014). Nitrogen disease hypothesis: Higher nutrient content of the plant material following nitrogen fertilization should promote disease. This is known for agricultural systems (Dordas 2008), but results from natural ecosystems vary (Mitchell *et al*. 2003; Veresoglou *et al*. 2013). Host dilution hypothesis: Many pathogens are dependent on the availability and density of host plants. At high plant diversity the abundance of each host plant is in average lower than in species poor communities (Civitello *et al*. 2015), which is suggested to be the underlying mechanism of observed negative diversity-disease relationships (Lau *et al*. 2008; Knops *et al*. 1999; Mitchell 2003; Mitchell *et al*. 2003).

## MATERIALS AND METHODS

### Experiment

We set up a large field experiment (PaNDiv Experiment) in the Swiss lowlands (mean annual temperature and precipitation 9.4±0.1°C, respectively 1021.62±31.89mm, MeteoSchweiz 2019) on a formerly extensively managed grassland in autumn 2015. The experiment consisted of 336 2m x 2m plots. We factorially manipulated plant species richness, plant functional composition (gradient of specific leaf area as a measure of growth strategy), nitrogen addition and foliar fungal pathogen exclusion. We used a set of 20 common grassland species spanning a large gradient of specific leaf area (SLA) to establish the experimental plant communities and divided them into fast (high SLA) and slow (Low SLA) growing (Table S2). The experimental communities contained either 1, 4, 8 or 20 species. Plots with 4 or 8 species could contain only slow, only fast or a mix of species, creating a large gradient in community mean SLA values. Monocultures spanned the full range in SLA values while plots with 20 species inevitably had an intermediate mean SLA. The communities had fully developed by late summer 2016. To maintain species compositions, the plots were weeded three times a year. Plots were mown once in the middle of June and once in August (for more details see Supplementary Methods and Pichon *et al*. 2019).

Each specific community composition received crossed nitrogen and fungicide treatments. Nitrogen (N) enrichment plots received 100 kgNha^-1^y^-1^, added once in April and once after the first mowing, in the form of urea. This is typical of fertilization experiments (Hautier *et al*. 2014) and medium – intensive farming (Bluethgen *et al*. 2012). Foliar fungal pathogen exclusion was done with fungicide (Score Profi by Syngenta Agro AG, 24.8% difenoconazole and Ortiva by Syngenta Agro GmbH, 22.8% azoxistrobin) applied four times during the growing season (0.2ml of Score Profi and 0.4ml of Ortiva mixed with 0.062l of water per treated plot each time). Plots without fungicide were sprayed with water. Difenoconazole interrupts the synthesis of ergosterol (IUPAC 2016), a fungal cell membrane component. If applied on top of the vegetation, it has no effect on soil (Dahmen, Staub 1992). Azoxistrobin blocks the cell respiration by inhibiting the proenzyme coenzyme Q which prevents the production of ATP. Studies have shown no phytotoxic effects of azoxistrobin (Sundravadana *et al*. 2007; Khalko *et al*. 2009) or difenoconazole (Nithyameenakshi *et al*. 2006). To account for potential soil heterogeneity across the study site, plots were arranged in four blocks. Each community composition was grown once per block and the nitrogen and fungicide treatments were assigned randomly to the communities in the blocks.

### Measurements

We measured plant aboveground **biomass** by harvesting two subplots of 0.1m^2^, 5cm above ground level, in mid-June and at the beginning of August 2018. Biomass was dried and weighed. Percentage cover of all sown plant species, plus weeds and bare ground, was visually estimated in the central square meter of the plots shortly before the biomass harvest (June and August). The sum of all estimates per plot could exceed 100% but here we analyse proportional abundances of each species. **Total plant cover** was calculated as 1- the proportion of bare ground. To describe the functional composition of the communities, we measured SLA (Garnier *et al*. 2001) on one leaf each from five plants, growing in the central square meter of all the monoculture plots (if possible, otherwise elsewhere in the plot) in June and in August, at the same time as we measured percentage cover. We then calculated several measures of plant functional composition. We calculated **the realized SLA**, i.e. the community weighted mean SLA per plot, using the percentage cover measurements and the mean SLA per monoculture as the baseline SLA for each species under a given treatment (the four combinations of nitrogen x fungicide). Because plant community composition can shift in abundance in response to the nitrogen and fungicide treatments, we also calculated the **shift in SLA** of the whole plant community relative to the **sown SLA** (mean SLA of all species sown in a community), by substracting the sown SLA from the realized SLA (see also Supplementary Methods).

Overall **fungal infection,** and infection with rusts, smuts, powdery mildews, downy mildews and leaf spots (see Rottstock *et al*. (2014), was assessed for each plant species in each plot, in July and in early October 2018. Ten randomly chosen individuals per plant species, growing in the central square meter of the plot (if possible, otherwise elsewhere in the plot), were screened for signs of infection and the percentage of infected individuals was recorded (see also Supplementary Methods). If there were less than 10 individuals in total, the percentage of infected individuals was calculated based on the observed number of individuals. Based on the species level infection, and the percentage cover of each plant species, we calculated an abundance weighted mean fungal infection per plot and season for total infection and infection by separate fungal groups (rusts, powdery and downy mildews and leaf spots). The smut fungi were excluded, because they were very rare (observed only eight times).

Further, we measured the **microclimate** (temperature and relative humidity logger iButton DS1923-F5, Maxim Integrated, USA) in each plot, in the center of one of the biomass subplots for a period of 2-3 days, with hourly measurements between 16.07.2018 and 13.08.2018. Due to a lack of data loggers we could only measure 28 plots at the same time (Table S3). Therefore, to account for differences in daily temperatures we subtracted the temperature and humidity measured in the plots from temperature and humidity measured at the same time in a nearby meteorological station in Zollikhofen (3.81 km away, MeteoSchweiz 2019).

### Analysis

Biomass, infection and trait data correlated well between the two time points when they were measured. For this reason, we used the total biomass (sum of the two harvests) and mean values of community shift SLA and fungal infection between the two time points.

We conducted two analyses to test for the causes and consequences of pathogen infection. We first analyzed the overall effects of fungicide on fungal infection and biomass production at the plot level, using linear mixed effect models, with fungicide as the independent variable and nitrogen addition, sown species diversity and realized SLA and all possible interactions as covariates. Block and species combination (84 levels) were included as random effects. We stepwise excluded non-significant terms from the model based on likelihood-ratio tests (Zuur 2009). We also ran separate models for each fungal group.

Secondly, we tested drivers and effects of quantitative levels of fungal pathogen infection in structural equation models (SEM, Figure S1, Table S1). As fungicide did not completely remove infection we fitted a multi-group SEM to test the drivers and effects of pathogen infection on control and fungicide plots separately (Grace 2006). All other treatment variables were also included in the SEM with direct effects on both fungal infection and biomass production. In the SEM we were therefore able to test for the effect of quantitative levels of pathogen infection and whether it varied with fungicide application. We included the deviation between plot and air humidity and temperature and plant cover, to account for indirect effects of the treatment variables through changes in microclimate. This considers potential impacts of the plant community on fungal infection, which elsewise would have likely influenced the path between fungal infection and biomass. We also incorporated an interaction between diversity and pathogen infection, which could affect plot biomass production, by constructing a dummy variable by multiplying the standardized values of fungal infection and species diversity (path 14 in Figure S1, Table S1).

We fitted a multi-group SEM, with the groups being the two levels of fungicide treatment, which allowed fungicide to interact with all the paths of the models. We checked whether each path and intercept differed significantly with fungicide, by comparing the AIC values of a fully unconstrained model, where all paths and intercepts were allowed to differ, with a model where a particular path was constrained to be equal between fungicide treatments. All paths that did not differ significantly were kept constrained (Table S2). We used the same SEM to analyze the separate fungal groups (rusts, powdery mildews, downy mildews and leaf spots).

To test for diversity effects through host dilution, we calculated host concentration effects for each plant species, as the relationship between host cover and infection. We fitted separate linear mixed effect models per plant species, nitrogen and fungicide treatment with block as a random effect. The slopes of these models were analyzed using another mixed effect model with nitrogen and fungicide as explanatory variables and species as a random effect. This allowed us to test whether nitrogen enrichment and fungicide alter any host dilution effects. All analyses were conducted in R (R Core Team 2018), using the package lme4 for linear mixed effects models (Bates *et al*. 2015) and lavaan for SEMs (Rosseel 2012).

## RESULTS

### Effects of fungicide application on fungal infection and plant biomass

Fungicide reduced fungal infection by 25.33% on average (Figure 2a). The fungicide was most effective in high SLA communities, especially at high species diversity and in the absence of nitrogen fertilization (Figure S4). Fungicide also increased plant biomass but only in plots with high SLA (Figure 2b). This agrees with the idea that fungicide was most effective in fast growing communities. Comparing the intercepts in the SEM between fungicide and non-fungicide plots showed similar results (Figure 4h).

**Figure 2.**
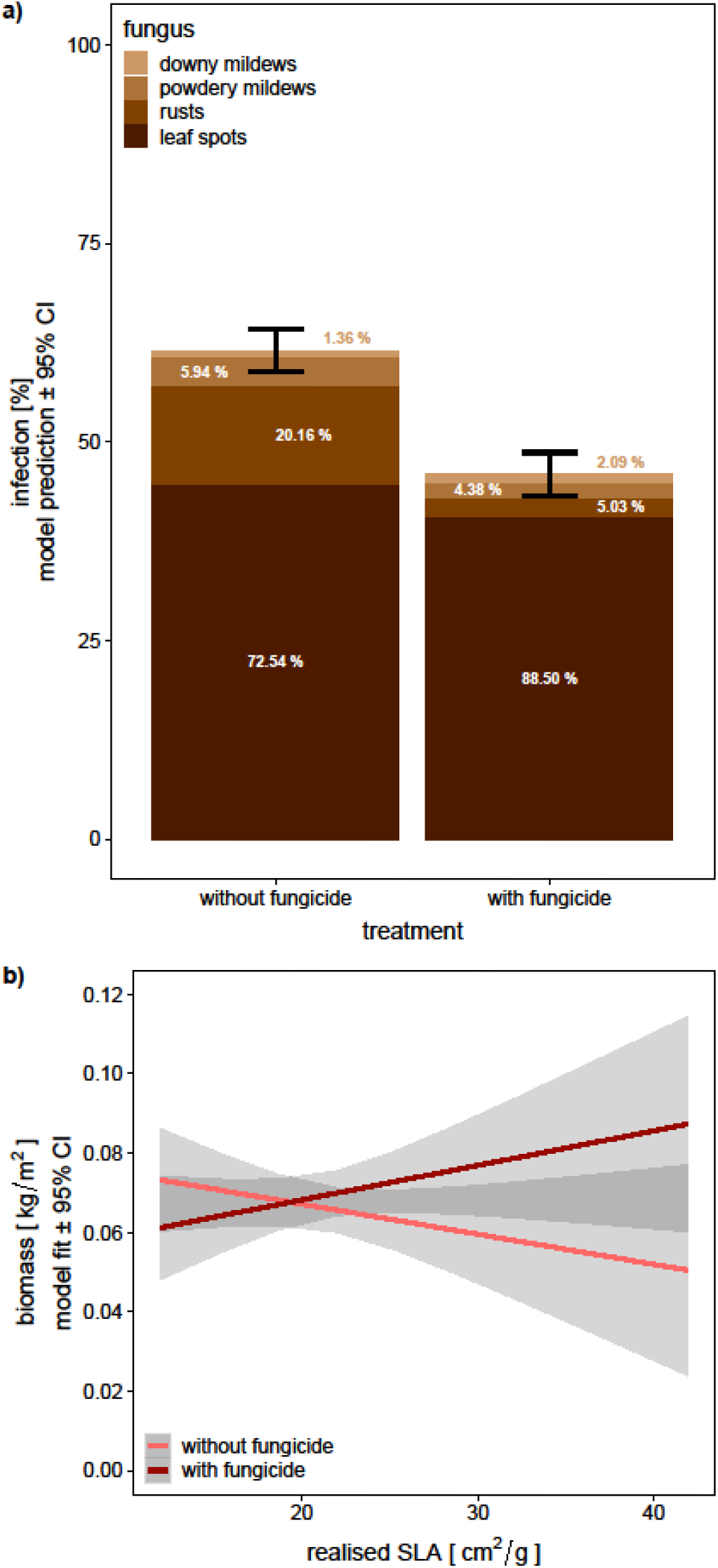
Selected results from the linear mixed effects models: model predictions and 95% confidence interval of a) impact of fungicide treatment on fungal infection and the contribution of single fungal groups to overall infection The numbers in the bars indicate the percentage contribution of each fungal group to the total infection. Main fungicide effects of the linear mixed effects models per fungal group: Fungicide reduced total infection from 61.50 ± 1.38 % to 45.92 ± 1.39 % (p < 0.001), leaf spots from 59.1 2 ± 1.58 % to 46.90 ± 1.57 % (p < 0.001), rusts from 16.43 ± 0.81 to 2.67 ± 0.81 % (p < 0.001) and powdery mildews from 4.85 ± 0.53 % to 2.32 ± 0.83 % (p < 0.001), while downy mildews were unaffected by fungicide (p = 0.623) and were generally very low (1.11 ± 0.39 %). b) Interactive effect of realised SLA and fungicide on biomass production. Plots dominated by fast-growing species produced less biomass than plots dominated by slow-growing species. Fungicide increased biomass production, but only in plots dominated by fast-growing species. Under fungicide treatment there was even an increase of biomass with increasing realised SLA. Estimates and CI were derived from the effects package (Fox 2003). The whole model results can be found in Table S4 and Table S5.

**Figure 3.**
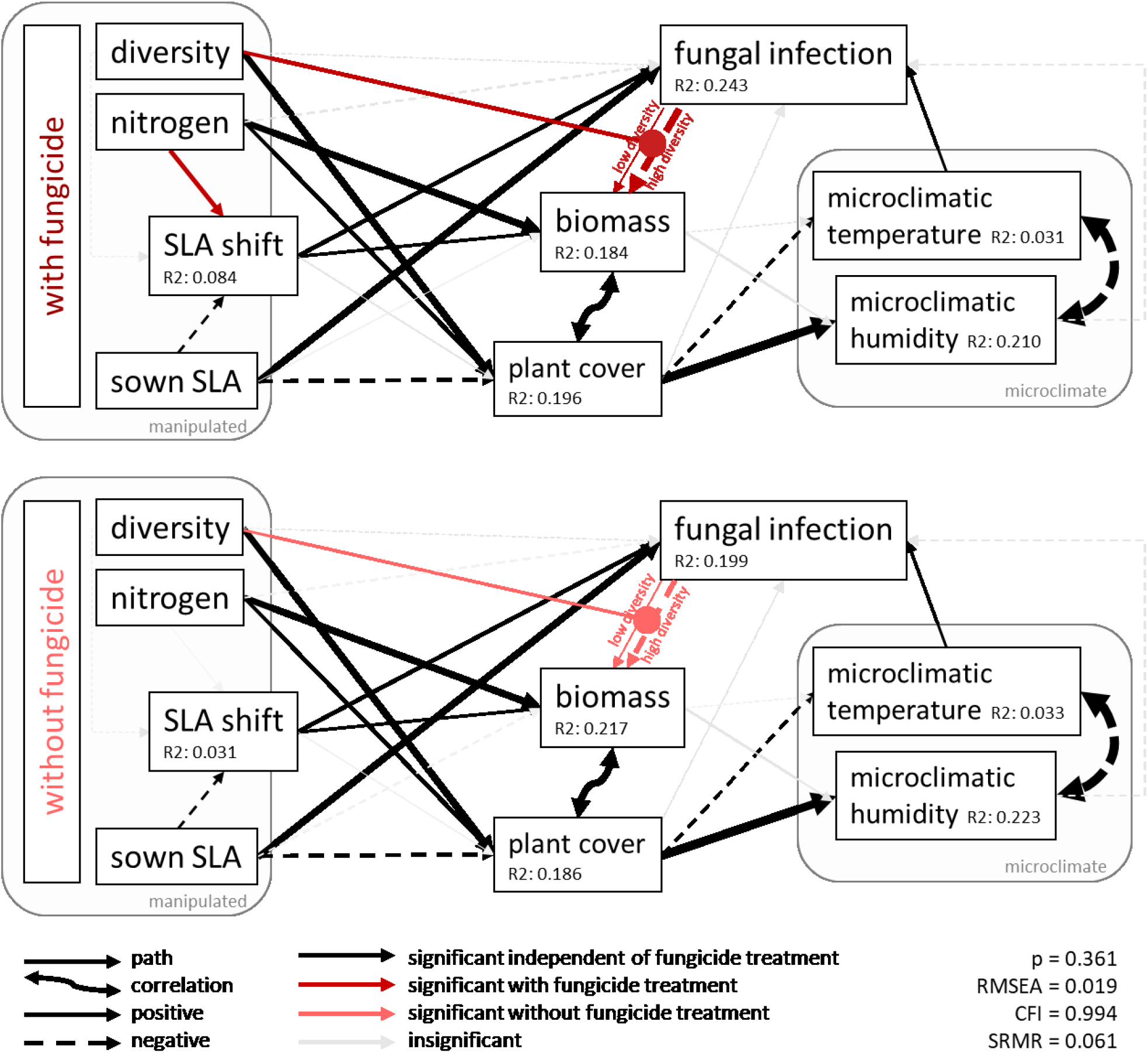
SEM: drivers and consequences of fungal infection. Dashed lines: negative effects. Solid lines: positive effects. Double headed arrows: correlations. Single headed arrows: paths. Black: significant constrained paths, red: significant unconstrained paths between fungicide (dark red) and no fungicide (light red). Light grey: not significant paths. Thickness: strength of the path/correlation.

**Figure 4.**
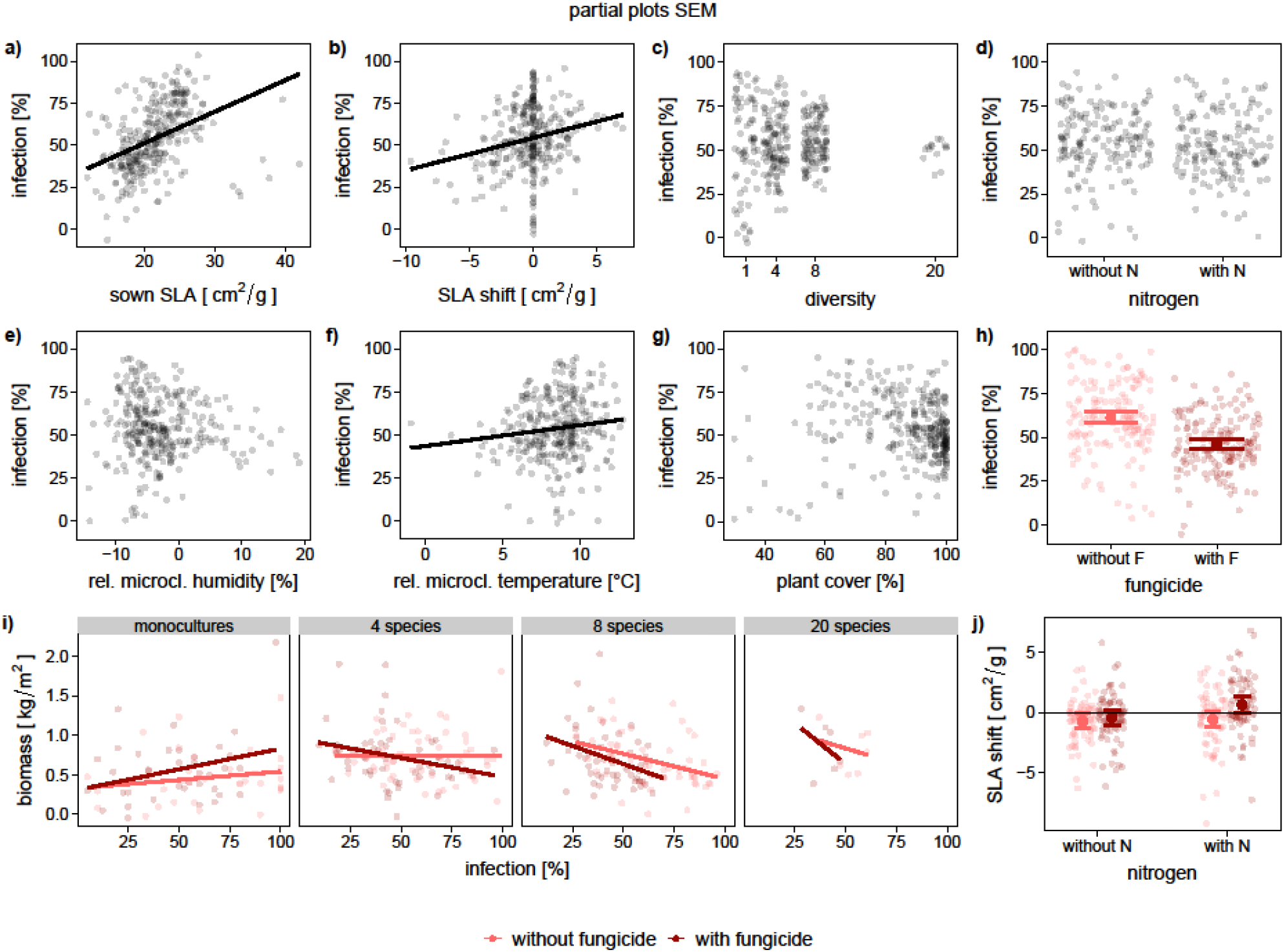
Partial plots of the SEM: impact of selected variables on fungal infection (a-h), biomass production (i) and SLA shift (j) after removing all effects of all the other variables which are not plotted. Effects on fungal infection of a) sown SLA (0.331, p<0.001), b) SLA shift (0.186, p<0.001), c) diversity (−0.030, p=0.551), d) nitrogen (−0.079, p=0.056), e) microclimatic humidity (−0.082, p=0.125), f) microclimatic temperature (0.121, p=0.012), g) plant cover (−0.061, p=0.227) and h) fungicide ±95% CI (−0.689, p<0.001) Interactive effects on biomass of i) fungicide, fungal infection and diversity and j) Interactive effects on SLA shift of nitrogen and fungicide estimate±95 & CI

### Drivers of infection

We then used SEM to look in more detail at the drivers of pathogen infection and its impacts on biomass (SEM: Figure 3; selected partial plots: Figure 4; path coefficients, significances, etc.: Table S8). The most important driver was functional composition, i.e. whether plant communities contained slow or fast growing plants. Both the sown SLA (Figure 4a) and the shift in SLA (Figure 4b) increased fungal infection. Communities with low sown SLA and a highly negative SLA shift, had lower infection than high SLA communities. Leaf spots, rusts and to some degree powdery mildews increased with increasing SLA, whereas downy mildews were unaffected (Figure S6).

Microclimate was also important and an increase in temperature increased fungal infection (Figure 4f). Humidity had no significant effect on fungal infection (Figure 4e). However, humidity and temperature were negatively correlated (Figure 3), which makes it hard to fully separate their effects. The impact of microclimate varied between fungal groups: rusts and leaf spots, the most abundant groups, increased with increasing temperature, while powdery and downy mildews were unaffected (Figure S6). Nitrogen and plant species diversity did not affect fungal infection directly (Figure 4c-d).

Several factors indirectly affected infection via changing microclimate. Temperature varied by 13.6°C between plots and was reduced by plant cover, but not by biomass. Plant cover was increased by plant diversity and nitrogen but reduced by sown SLA. Therefore, in addition to its positive direct effect, sown SLA also had a positive indirect effect on fungal infection, but this indirect path was non-significant overall. Species diversity and nitrogen enrichment indirectly decreased infection by increasing plant cover and reducing temperature, but again the indirect effects were not significant overall.

The absence of a direct diversity effect on fungal infection cannot be explained by an absence of host concentration effects, as on average plant species cover was positively related to species-specific infection, suggesting additional mechanisms such as spillover or additionally the presence of density independent pathogens. The application of fungicide removed host concentration effects (Figure S7).

### Impact of fungal infection

In the SEM (SEM: Figure 3; selected partial plots: Figure 4; path coefficients, significances, etc.: Table S8), fungal infection also affected plant biomass production, however this depended on plant diversity (Figure 4i): in species rich communities fungal infection was negatively related to plant biomass, indicating that fungi had strong impacts on biomass, whereas in monocultures, fungal infection was even weakly positively related to biomass (Figure 4i). Adding fungicide increased the effect of diversity on the disease-productivity relationship, which means a stronger negative correlation at high diversity and a stronger positive correlation between infection and biomass in monocultures (Figure 4i). The SEMs per fungal group revealed that the leaf spots and to some degree the rusts drove the negative relationship between infection and biomass (Figure S6). Powdery mildew had no impact on biomass production, while the downy mildews even increased biomass. The downy mildew and rust models did not fit well (both p<0.001) but the fit was good for the leaf spots (p = 0.268) and adequate for the powdery mildews (p = 0.088).

Biomass was also affected by several other factors. Nitrogen enrichment increased biomass production independently of the fungicide treatment, a shift in SLA towards faster growing species increased biomass production, while the effects of sown SLA on biomass depended on the fungicide treatment.

Fungicide also altered the SLA shift in the experimental plant communities to favor faster growing species (Figure 4j), but there was a lot of unexplained variation in SLA shift (R^2^=0.084 under fungicide and R^2^=0.031 under no fungicide treatment). The effect of fungicide on the SLA shift was amplified by nitrogen enrichment so that plots with nitrogen added and pathogens reduced shifted towards dominance by faster growing species (Figure 4j).

## DISCUSSION

### Growth-defense trade-off

We found strong support for the growth-defense trade-off hypothesis as the key driver of pathogen abundance and impact. Plant communities dominated by fast growing species had increased infection, and fungicide was most effective at reducing fungal infection in high SLA communities. There is an inherent trade-off between plant growth and the production of certain defense compounds (Huot *et al*. 2014), and species which are at the fast end of the leaf economics spectrum have been shown to have lower structural and chemical defenses (Mason *et al*. 2016; Coley 1988) and higher tissue nutrient levels (Wright *et al*. 2004). Both could explain the increased pathogen attack on fast growing species, however, the absence of support for the nitrogen disease hypothesis, see below, may indicate that changes in defenses are more important. Our results show that growth-defense trade-offs are not only a major predictor of herbivory (e.g. Lind et al.) and pathogen attack on individual plant species but also scale up to be the key driver of community level pathogen infection.

Fungal pathogen impact was also mostly determined by growth-defense trade-offs. Fungicide allowed fast growing species to increase in abundance, especially under nitrogen. This is in line with findings that plants originating from nutrient rich habitats benefitted most from enemy release (Blumenthal *et al*. 2009, but see e.g. Heckman *et al*. 2017). Fast growing species are expected to be good competitors in nutrient rich environments (Wright *et al*. 2004; Poorter *et al*. 2009), but our results suggest that pathogens reduce their competitive advantage. Pathogens may therefore equalize competitive abilities and promote diversity in nitrogen rich conditions. In nutrient poor habitats, slow growing plants are expected to be more competitive and in such an environment, pathogens might reduce diversity by excluding faster growing species. Previous studies have shown pathogens can alter the outcome of plant competition (Paul 1989; Ridenour & Callaway 2003) and change plant community composition (Allan et al. 2010). Our results suggest that the growth strategy of plants is the key predictor of plant community responses to pathogens, and that pathogens promote slow growing species. Over time, this would be expected to reduce pathogen abundance and therefore impact. Such feedbacks could cause temporal dynamics between plant community composition and fungal infection, which could only be tested with long term data on fungal infection and plant functional composition.

### Nitrogen disease

We did not find support for the nitrogen disease hypothesis. Nitrogen can increase disease in crops but findings from grasslands are contradictory, with some studies finding support (Mitchell *et al*. 2003), but others not (Lau *et al*. 2008). Compared to agricultural systems, grassland plants could evolve increased disease resistance with nitrogen fertilization (Snaydon & Davies 1972), which might offset any benefits the pathogens would derive from higher plant nutrient contents. In addition, plant community composition changes with nitrogen enrichment. Mitchell *et al*. (2003) did not control for changes in composition but showed that the “disease proneness” of the plants was an important driver of infection. Liu *et al*. (2018) showed that nitrogen addition favors disease prone species (but see Welsh *et al*. 2016). However, these studies did not explain what drives disease resistance and could not separate compositional change effects from direct effects of nitrogen. Our results indicate that trade-offs linked to the leaf-economics spectrum are likely the underlying mechanism and that an increase in fast growing species is responsible for an increase in infection with nitrogen. Further, nitrogen enrichment can increase humidity and decrease temperature through increased shading in denser vegetation. In the dry summer of 2018, N fertilisation may have decreased water and temperature stress and made the plants more resistant to fungal infection, which would explain why we found a negative indirect effect of nitrogen enrichment on infection. This all suggests that the direct effect of nitrogen on community infection in grasslands is weak to non-existent. Nitrogen enrichment rather drives infection through indirect effects of community shift and changes in microclimatic conditions.

### Impact of plant diversity

Plant diversity did not affect fungal infection in our study, apart from a small indirect effect through microclimate. This is contrary to most other studies, which found that an increase in diversity leads to a decrease in infection (e.g. Mitchell *et al*. 2002; Liu *et al*. 2016; Rottstock *et al*. 2014; but see Halliday *et al*. 2017). We expected that host abundance would be diluted at high plant diversity and that this would reduce infection. However, while infection on individual plant species was lower when the plants were rarer (at least when pathogens were not suppressed by fungicide), this did not lead to a negative diversity-infection relationship for the community. Other diversity related mechanisms may have counteracted this relationship. Several other studies reported unexplained effects of diversity on fungal infection, in addition to host dilution, and different plant species and diseases varied in their response to diversity (Rottstock *et al*. 2014; Mitchell *et al*. 2002; Knops *et al*. 1999). One mechanism by which diversity can counteract dilution effects is increased spillover of generalist pathogens at high diversity or an increase of density independent pathogens such as vector-transmitted ones (Power & Mitchell 2004; Halliday *et al*. 2017). Another possibility is that diverse communities become dominated by susceptible species, limiting host dilution effects. Both mechanisms might explain why plant diversity did not affect fungal infection in our study.

Interestingly, our results suggest that the impact of fungal pathogens on biomass production was higher in species rich plant communities. Even though plant diversity did not alter overall pathogen infection, it could still have altered fungal community composition or diversity and might have led to more aggressive fungi at high plant diversity or reduced pathogen tolerance of the plants. However, we did not find that diversity altered the abundance of our four fungal guilds. It is therefore also possible that the ability of the plants to deal with infection varies with diversity. A higher pathogen pressure in species poor communities might select for better defended plant genotypes, leading over time to reduced pathogen impact in monocultures. Results from the Jena Experiment support this idea and show that plants in monocultures have evolved to be more resistant against belowground pathogens (Zuppinger-Dingley *et al.)* and aboveground fungi (Hahl *et al*. 2017). To better predict variation in pathogen impact in plant communities we may need to consider pathogen community composition and host genetics.

### Climatic stress

Temperature also affected fungal infection - leaf spots and rusts both benefitted from an increase in temperature in the vegetation. Other studies also indicate that higher temperatures increase pathogen infection (Liu *et al*. 2016) and that different fungal groups vary in their responses. Powdery mildews can increase with temperature, while rusts show more variables responses (Gullino *et al*. 2018; Helfer 2014). Longer periods of 100% humidity lead to water condensation, which has been shown to increase infection (Burdon 1991; Sun *et al*. 2017). However, we found no effect of humidity on infection, after correcting for temperature. The summer 2018 was extraordinarily hot and dry, with mean July temperatures 1.6°C above the average of the last 30 years and precipitation 18.81% lower (MeteoSchweiz 2019), which likely resulted in intensive drought and heat stress for the plants. Drought stress can increase fungal diseases in trees (Desprez-Loustau *et al*. 2006) and increase the negative effects of pathogens on competitive ability (Paul & Ayres 1987). The microclimate itself was driven by plant cover, which was determined by plant diversity, nitrogen and functional composition. These variables indirectly (but weakly) influenced fungal infection through a change in the microclimate. Changes in vegetation microclimate may therefore play an important role in affecting plant community resistance to disease under extreme weather conditions.

### Impact of fungicide and infection intensity on plant biomass

In our study we used two approaches to assess the impact of fungal pathogens: exclusion with fungicide and SEMs testing the effect of infection intensity on plant biomass production. Fungicide application increased plant biomass but only in plots dominated by fast growing plants, which suggests that fast growing plants are not entirely tolerant. The magnitude of biomass reduction was lower than in other studies (Allan *et al*. 2010; Seabloom *et al*. 2017), perhaps because, unlike in the other studies, we mowed the field regularly, preventing the build-up of large pathogen populations over the season. Our analysis relating infection and biomass suggested that the negative impact of fungal infection on biomass in high diversity plots was amplified by fungicide, even though fungicide generally decreased infection. Fungicide shifted the functional composition of the fungal community by mainly removing the rather specialized rusts and powdery mildews (Klenke 2015) and it removed host concentration effects, which also suggests a shift from specialists towards more generalist pathogens (Bever *et al*. 2015). Fungicide may therefore have selected for more aggressive, generalist pathogens, which would also explain its small overall effect on biomass production. These results suggest that a shift in pathogen community composition could be a major driver of pathogen impact. Many studies assess the impact of fungal infection on ecosystem functioning by comparing plant biomass in fungicide and non-fungicide plots (Mitchell 2003; Allan *et al*. 2010; Seabloom *et al*. 2017; Heckman *et al*. 2017). Our results show the importance of complementing these experiments with measures of infection severity and pathogen community composition. To increase our mechanistic understanding of the role of pathogens in affecting ecosystem functioning it is crucial to combine both approaches.

One alternative explanation for the altered impact of fungal infection on biomass under fungicide treatment might be non-target effects of the fungicide. However, studies show that the fungicides used here do not have phytotoxic effects when they are used in the recommended concentrations (Sundravadana *et al*. 2007; Khalko *et al*. 2009; Nithyameenakshi *et al*. 2006). Fungicides might also reduce beneficial fungi, like mycorrhiza belowground, or other mutualistic leaf-endophytes (Fokkema & Nooij 1981; Henriksen & Elen 2005). However, root samples of a subset of the experimental plant species showed no difference in mycorrhizal colonization between plants from fungicide and non-fungicide plots in 2017 (data not shown) and a loss of mutualists would be expected to reduce biomass production with fungicide application. This suggests that while non-target effects cannot completely be excluded, they are unlikely to be the key driver of our results.

## Conclusions

We found strong support for growth-defense trade-off as a main driver of fungal infection. Fungal infection had an impact on biomass production, but this impact was context dependent, with greatest biomass loss due to pathogens in species rich communities receiving fungicide treatment. Fungicide altered the complex plant-pathogen interactions, beyond just removing pathogens, probably by removing certain fungi more efficiently than others. Fungicide may therefore have a wider range of effects in ecosystems than previously considered. This is both a challenge and an opportunity for studies using fungicide treatments.

## AUTHORS’ CONTRIBUTIONS

SC, NP and EA designed and set up the PaNDiv experiment. NP and SC collected the data. SC analyzed the data and wrote the manuscript with substantial input from EA, NP and AK.

## AKNOWLEDGEMENTS

We thank Hugo Vincent and the whole PaNDiv team and helpers, without whom maintaining such a large experiment would not have been possible. Thanks to Nadia Maroufi and Tosca Manall for their feedback on the manuscript and to Fletcher Halliday for the friendly review. This study was supported by funding of the Swiss National Science Foundation.

## SUPPLEMENTARY

### Supplementary Methods

#### Experiment

We set up a large field experiment (PaNDiv Experiment) in the Swiss lowlands, close to the city of Bern (mean annual temperature and precipitation 9.4±0.1°C, respectively 1021.62±31.89mm, MeteoSchweiz 2019). The grassland contains a species composition typical for a nutrient rich, rather dry, grazed grassland (Delarze 2015). We cleared an area of 3145m^2^ (85m x 37m) of all vegetation in autumn 2015 and sowed our experimental plant communities. Some species were resown in spring 2016, because of poor establishment. The experiment consisted of 336 2m x 2m plots, separated by a 1m path sown with a grass seed mixture consisting of *Lolium perenne* and *Poa pratensis* (UFA-Regeneration Highspeed) and mown regularly during the growing season.

We factorially manipulated plant species richness, plant functional composition, nitrogen addition and foliar fungal pathogen exclusion. We used a set of 20 common grassland species to establish the experimental plant communities (Table S2). Half of the species were classified as fast, half as slow-growing based on specific leaf area (SLA) and leaf nitrogen content (Figure S2), which are traits indicative of the leaf economics spectrum (Reich & Cornelissen 2014; Wright *et al*. 2004). We did not include legumes in the species pool, because most legumes are adapted to low nitrogen levels and could therefore have been only included in the slow species pool only, making the species pools phylogenetically biased. The experimental communities contained either 1, 4, 8 or 20 species. Plots with 4 or 8 species could have either only slow-growing species, only fast-growing species or a mixture of both, which created a large gradient in community weighted mean traits. We grew monocultures of all species, which were either fast- or slow-growing, and the plots containing all 20 species inevitably had mixed functional compositions. The species for 4 and 8 species communities were chosen randomly from their respective species pools. To maintain species compositions, the plots were weeded three times a year. Plots were mown once in the middle of June and once in August, close to the dates when the farmers usually mow their extensive meadows (for more details see Pichon *et al*. (2019)).

### Conceptual SEM

**Figure S1.**
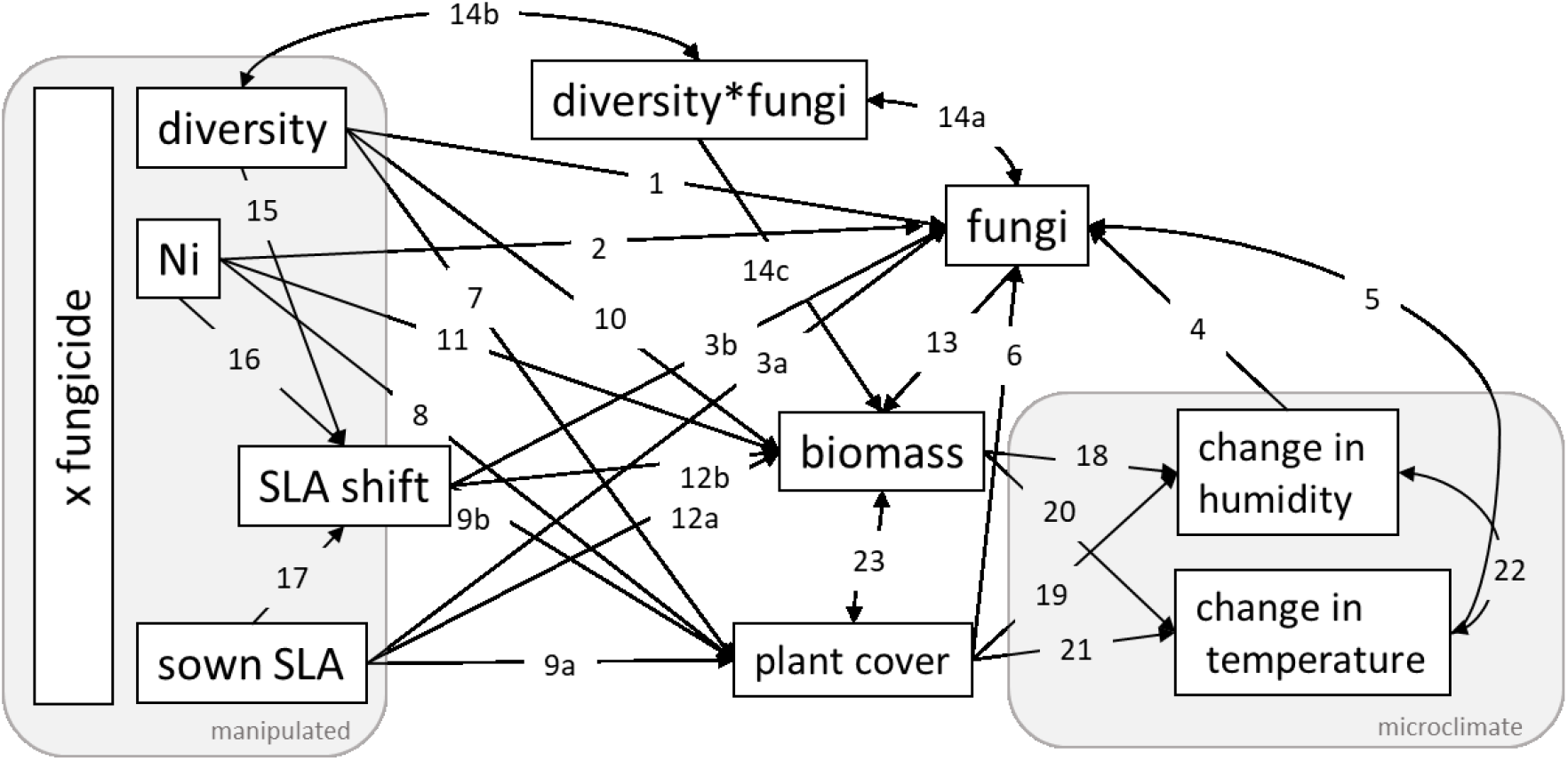
The full SEM that we tested. We tested all paths from the manipulated variables (left box) to the measured variables. The numbered paths are explained in Table S1.

**Table S1.** Hypothesized mechanism driving fungal infection and biomass production in the SEM model (Figure S1)

| Path | Hypothesized mechanism | Reference |
| --- | --- | --- |
| 1 | (?/-) mostly negative diversity effect on infection in grasslands, through host dilution | Rottstock *et al.* (2014), Mitchell *et al.* (2003), Mitchell *et al.* (2002) |
| 2 | (+) nitrogen disease | Dordas (2008) |
| 3 | (+) growth defense trade-off | Wright *et al.* (2004) |
| 4 | (+) high humidity is often beneficial for fungal pathogen growth and sporulation | Bregaglio *et al.* (2013), Chen *et al.* (2014), Sun *et al.* (2017), Bradley *et al.* (2003) |
| 5 | (+) an increase in temperature can increase fungal infection | Liu *et al.* (2016), Roy *et al.* (2004) |
| 6 | (?/+) spillover of generalist fungi because of higher density of potential hosts | Power & Mitchell (2004), Parker *et al.* (2015) |
| 7,10 | (+) positive diversity-productivity relationship | Tilman *et al.* (2001) |
| 8,11 | (+) nitrogen increases productivity due to nutrient limitation | Whitehead (1970), DiTommaso & Aarssen (1989), Fay *et al.* (2015) |
| 9,12 | (+) high growth rate at high SLA | Reich *et al.* (1992), Lavorel & Grigulis (2012) |
| 13 | (-) consumption of biomass through fungal pathogens | Allan *et al.* (2010), Seabloom *et al.* (2017) |
| 14 | (±) different selection pressure (due to host dilution), pathogen community composition shift | Laine (2006), Roy *et al.* (2000) |
| 15 | (±) increase in competition for light, sampling effect | Bachmann *et al.* (2018) |
| 16 | (+) nitrogen enrichment favors the abundance of fast growing plants | Lavorel & Grigulis (2012), Liu *et al.* (2018), De Vries *et al.* (2012) |
| 17 | (±) likely the communities with extremely high/low sown SLA have the biggest negative/positive shifts towards intermediate SLA |  |
| 18, 19 | (+) more plant transpiration in denser vegetation, shelter against wind that could remove humid air | Procházka *et al.* (2011) |
| 20, 21 | (-) more shading | Procházka *et al.* (2011) |

**Table S2.** PaNDiv experimental species.

| Species | resource economics | group |
| --- | --- | --- |
| *Dactylis glomerata* | fast | grass |
| *Holcus lanatus* | fast | grass |
| *Lolium perenne* | fast | grass |
| *Poa trivialis* | fast | grass |
| *Anthriscus sylvestris* | fast | herb |
| *Crepis biennis* | fast | herb |
| *Galium album* | fast | herb |
| *Heracleum sphondylium* | fast | herb |
| *Rumex acetosa* | fast | herb |
| *Taraxacum officinale* | fast | herb |
| *Anthoxanthum odoratum* | slow | grass |
| *Bromus erectus* | slow | grass |
| *Festuca rubra* | slow | grass |
| *Helictotrichon pubescens* | slow | grass |
| *Achillea millefolium* | slow | herb |
| *Centaurea jacea* | slow | herb |
| *Daucus carota* | slow | herb |
| *Plantago media* | slow | herb |
| *Prunella grandiflora* | slow | herb |
| *Salvia pratensis* | slow | herb |

**Figure S2.**
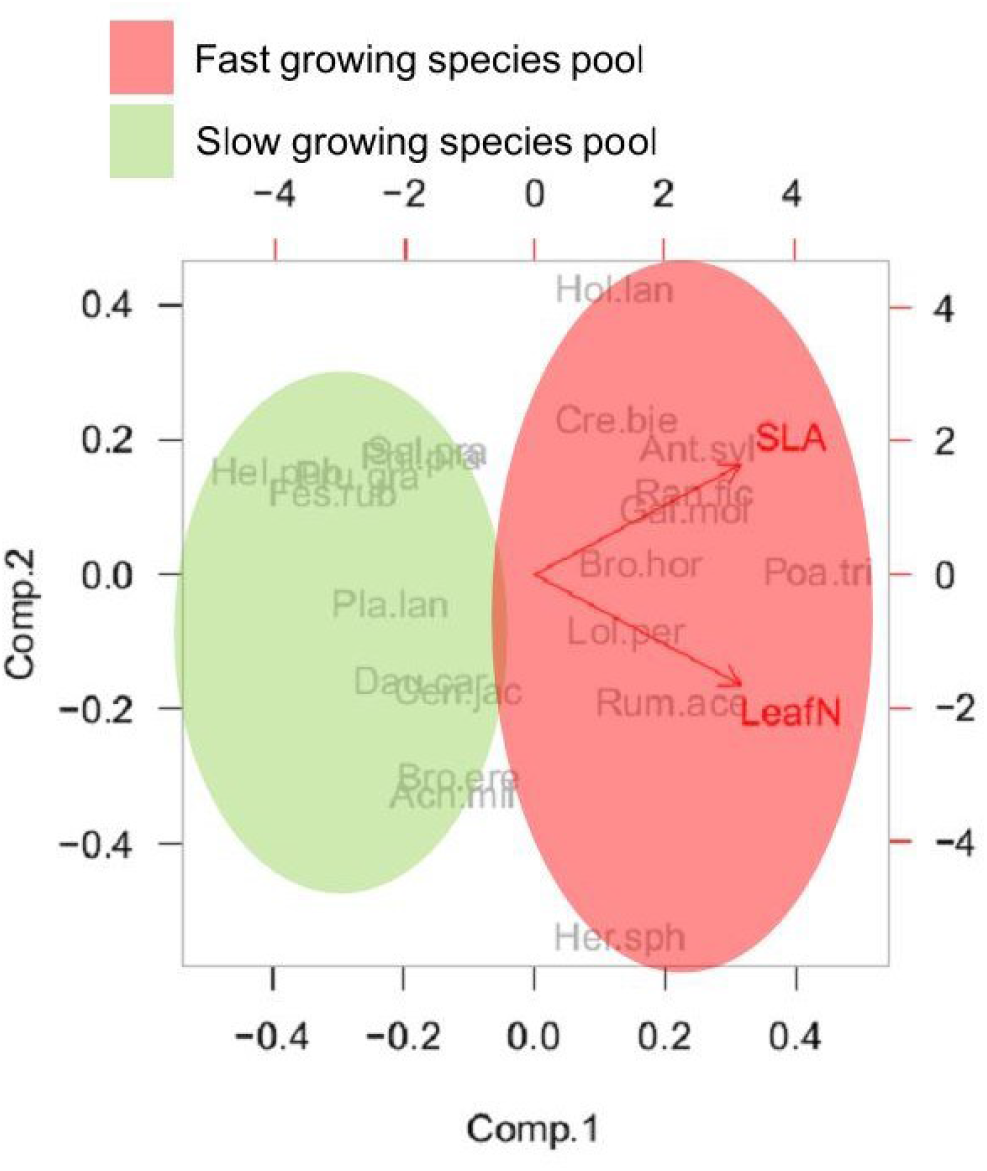
PCA categorizing the experimental species as fast and slow growing based on their values of SLA and leaf nitrogen

#### Measurements

##### Measures of SLA

By including the sown SLA and the SLA shift per plot, calculated based on the monoculture measurements with the corresponding nitrogen and fungicide treatment we accounted for abundance shifts and for plastic shifts following nitrogen and fungicide treatments, but not plastic shifts as a response to diversity. The latter, however is not significant compared to the plastic shifts as a response to nitrogen and fungicide (data not shown).

##### Infection

Measuring the % of infected individuals is different from many studies, which measure infeaction as damaged leaf area (e.g. Mitchell (2003), Halliday *et al*. (2017)), but likely more suitable to compare different fungal groups, as some (e.g. powdery mildews) mainly grow on the leaf, while others (e.g. rusts) mainly grow in the leaves, which makes a big part of the infection invisible (Klenke 2015). Percent leaf area damaged and percent infected individuals are log-correlated (Figure S3) and are therefore not fundamentally different from each other.

**Figure S3.**
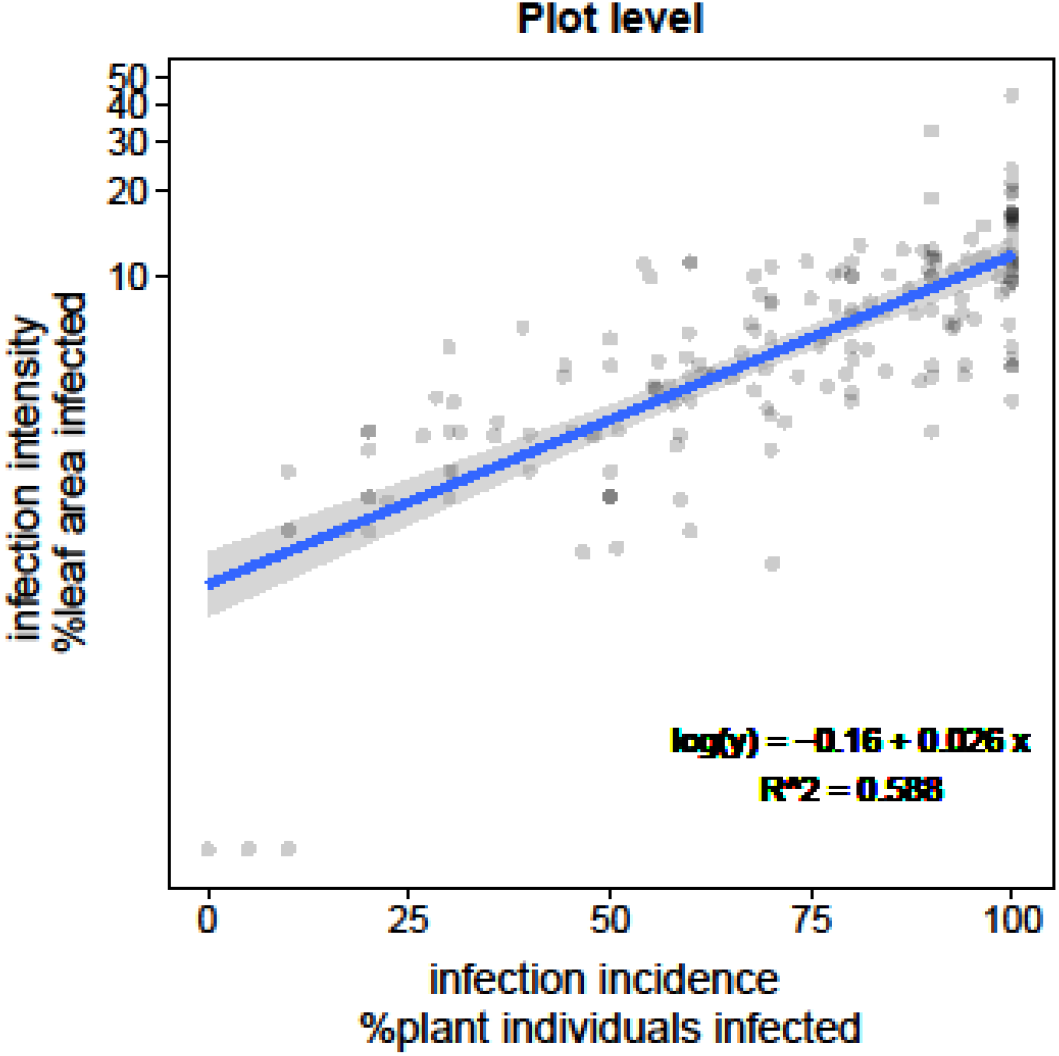
Correlation between community weighted mean of infection intensity based on % leaf area infected and community weighted mean of infection incidence based on % infected individuals. Data from fall 2018, as damaged leaf area was only assessed in fall 2018.

##### Microclimate

**Table S3.**
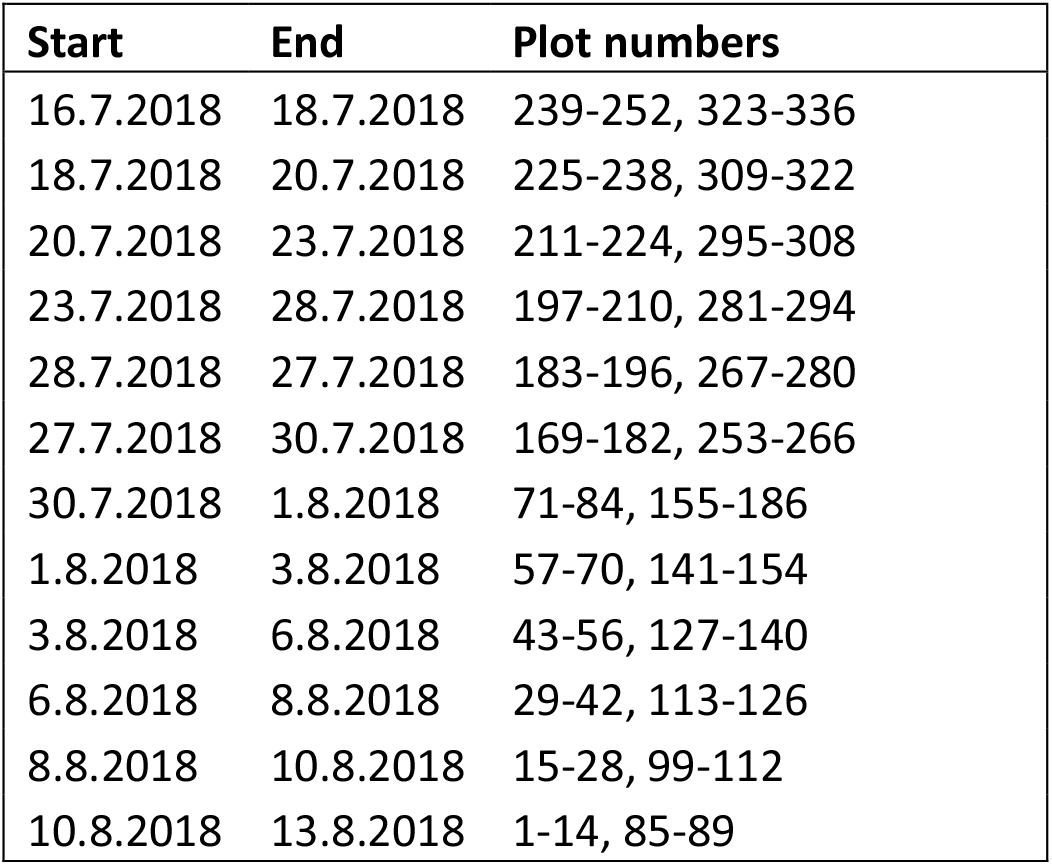
Dates when humidity and temperature loggers were placed in which plots

## Supplementary Analyses

**Table S4.** fixed effects of the fungi lmer

| Fixed Effects | Estimate | SE | t value | Chi² | p-value |
| --- | --- | --- | --- | --- | --- |
| Intercept | 0.42213 | 0.09771 | 4.32 |  | marginal |
| Nitrogen | -0.10653 | 0.08237 | -1.293 |  | marginal |
| Fungicide | -0.78739 | 0.08393 | -9.382 |  | marginal |
| Species Diversity | -0.02767 | 0.08368 | -0.331 |  | marginal |
| Realized SLA | 0.48125 | 0.08995 | 5.35 |  | marginal |
| Nitrogen x Fungicide | 0.08203 | 0.12109 | 0.677 |  | marginal |
| Fungicide x Species Diversity | -0.06974 | 0.05908 | -1.18 |  | marginal |
| Nitrogen x Realized SLA | -0.13115 | 0.08343 | -1.572 |  | marginal |
| Fungicide x Realized SLA | -0.40054 | 0.09638 | -4.156 |  | marginal |
| Species Diversity x Realized SLA | 0.23772 | 0.08665 | 2.743 |  | marginal |
| Nitrogen x Fungicide x Realized SLA | 0.36719 | 0.12227 | 3.003 | 9.187 | 0.002 |
| Species Diversity x Fungicide x Realized SLA | -0.223 | 0.09118 | -2.446 | 6.128 | 0.013 |
| Random Effects | Variance | SD |  |  |  |
| Composition | 0.4232 | 0.6505 |  |  |  |
| Block | 0.0031 | 0.0560 |  |  |  |

**Figure S4.**
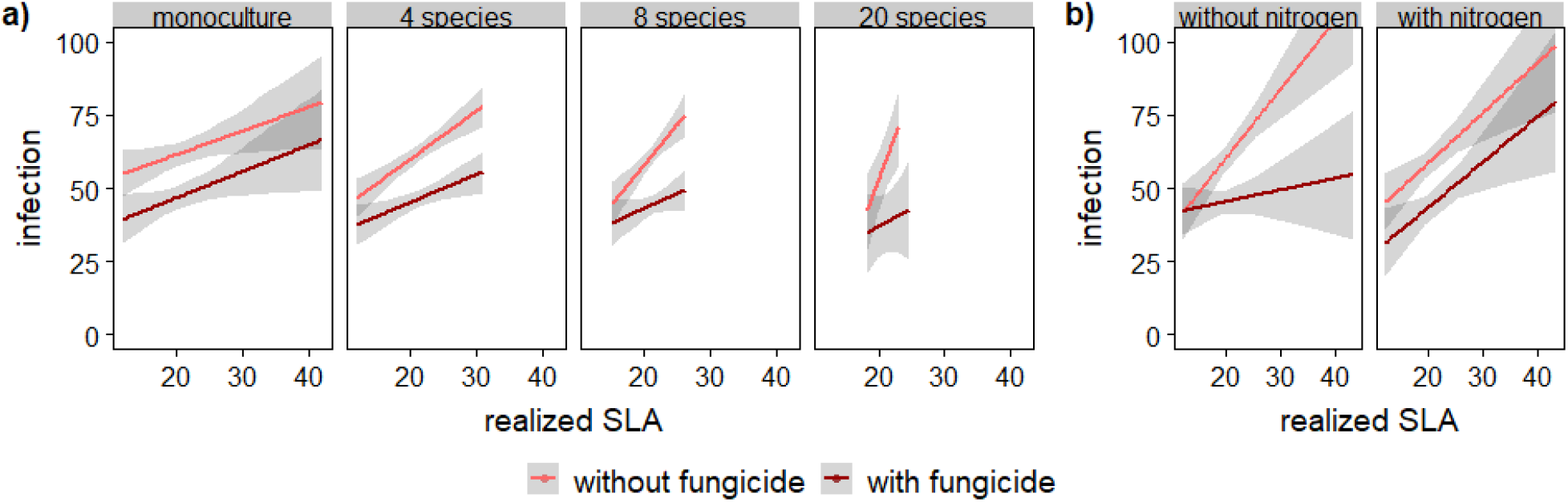
model predictions of lmer for fungal infection of all significant interactions terms (obtained from the effect package in r (Fox 2003)). Fungicide had significant interactions with a) fungicide, realized SLA and plant species diversity in explaining fungal infection, and with b) SLA as well as nitrogen in explaining biomass production. Fungicide reduced fungal infection on average (t-value= −11.942, infection without fungicide: 61.50 ± 1.38 %, infection with fungicide: 45.92 ± 1.39 %). Estimates and CI were derived from the effects package (Fox 2003)

**Table S5.** fixed effects of the biomass lmer

| Fixed Effects | Estimate | SE | t value | Chi<sup>2</sup> | p-value |
| --- | --- | --- | --- | --- | --- |
| Intercept | -0.3259 | 0.1393 | -2.3400 |  | marginal |
| Nitrogen | 0.5828 | 0.0881 | 6.6120 | 41.49 | <0.001 |
| Fungicide | 0.0779 | 0.0864 | 0.9019 |  | marginal |
| Realized SLA | -0.0865 | 0.0768 | -1.1266 |  | marginal |
| Species Diversity | 0.1374 | 0.0688 | 1.9966 | 3.999 | 0.046 |
| Fungicide x Realized SLA | 0.1859 | 0.0879 | 2.1138 | 4.518 | 0.034 |
| Random Effects | Variance | SD |  |  |  |
| Composition | 0.2340 | 0.4837 |  |  |  |
| Block | 0.0330 | 0.1817 |  |  |  |

**Figure S5.**
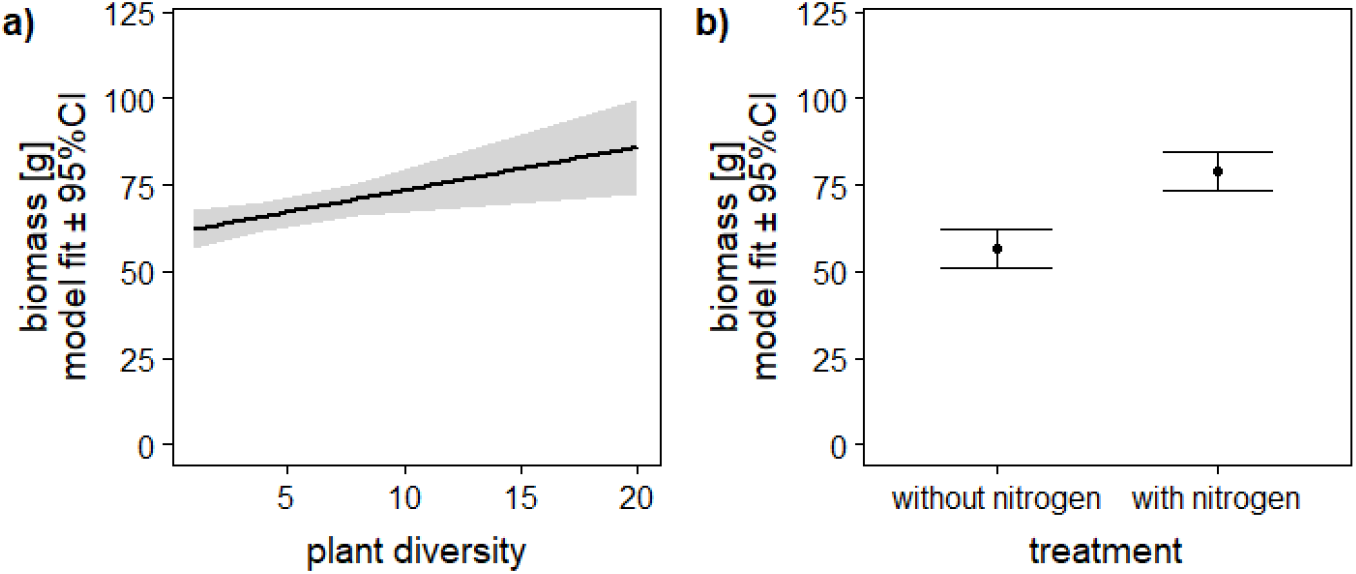
model predictions of the biomass lmer for a) plant diversity and b) nitrogen treatment (obtained from the effect package in r (Fox 2003)).

**Table S6.**
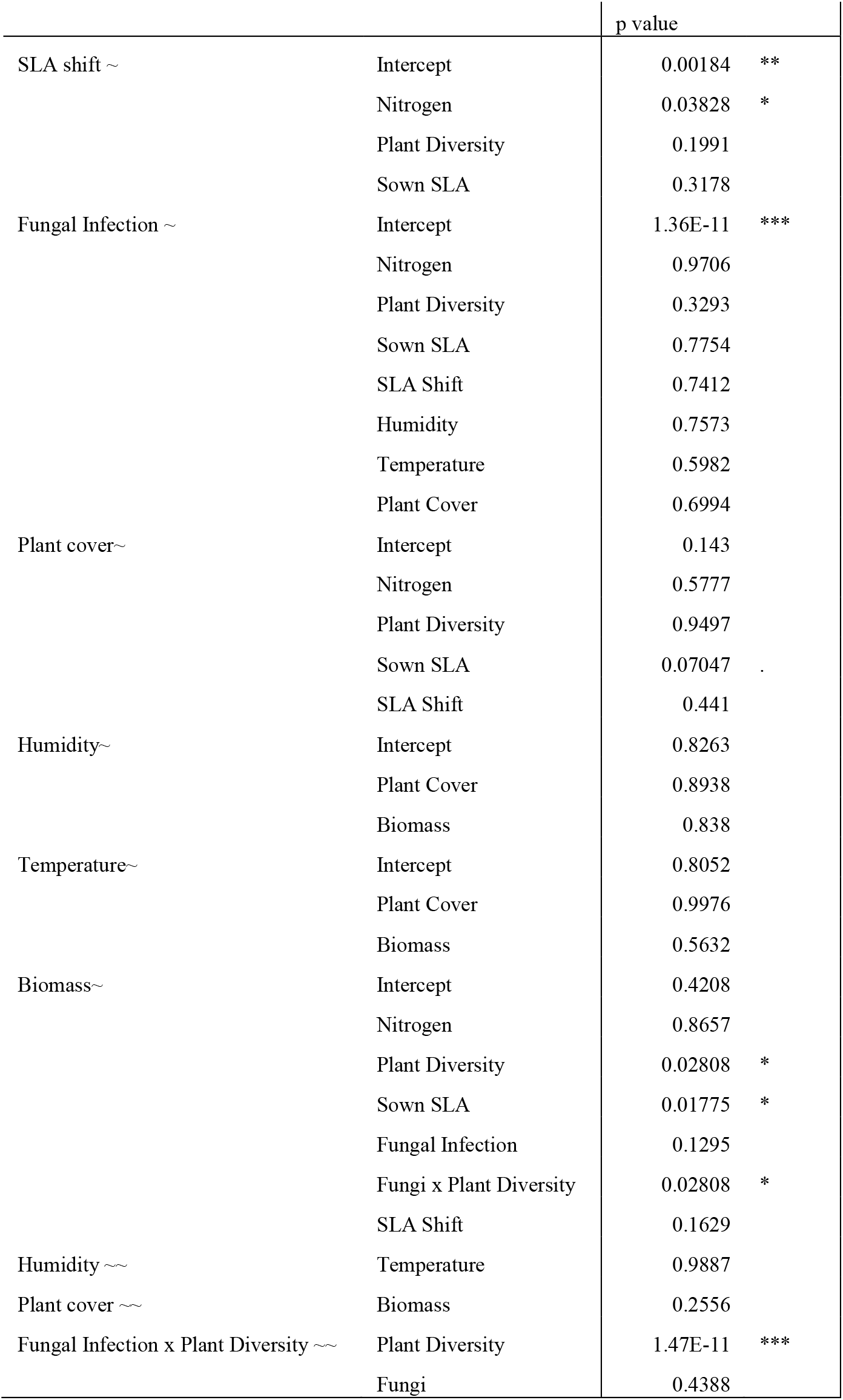
Results from the SEM model constrain critera, where we tested whether paths and intercepts differed significantly with fungicide, by comparing the AIC values of a fully unconstrained model with a model where a particular paths was constrained to be equal between fungicide treatments. Note that we did not constrain the path between biomass and fungal infection, even though it does not significantly differ between treatments. Fungal infection is part of the interaction term, which cannot be constrained, we therefore did not constrain any paths that are part of this interaction.

|  |  | p value |  |
| --- | --- | --- | --- |
| SLA shift ~ | Intercept | 0.00184 | ** |
|  | Nitrogen | 0.03828 | * |
|  | Plant Diversity | 0.1991 |  |
|  | Sown SLA | 0.3178 |  |
| Fungal Infection ~ | Intercept | 1.36E-11 | *** |
|  | Nitrogen | 0.9706 |  |
|  | Plant Diversity | 0.3293 |  |
|  | Sown SLA | 0.7754 |  |
|  | SLA Shift | 0.7412 |  |
|  | Humidity | 0.7573 |  |
|  | Temperature | 0.5982 |  |
|  | Plant Cover | 0.6994 |  |
| Plant cover~ | Intercept | 0.143 |  |
|  | Nitrogen | 0.5777 |  |
|  | Plant Diversity | 0.9497 |  |
|  | Sown SLA | 0.07047 | . |
|  | SLA Shift | 0.441 |  |
| Humidity~ | Intercept | 0.8263 |  |
|  | Plant Cover | 0.8938 |  |
|  | Biomass | 0.838 |  |
| Temperature~ | Intercept | 0.8052 |  |
|  | Plant Cover | 0.9976 |  |
|  | Biomass | 0.5632 |  |
| Biomass~ | Intercept | 0.4208 |  |
|  | Nitrogen | 0.8657 |  |
|  | Plant Diversity | 0.02808 | * |
|  | Sown SLA | 0.01775 | * |
|  | Fungal Infection | 0.1295 |  |
|  | Fungi x Plant Diversity | 0.02808 | * |
|  | SLA Shift | 0.1629 |  |
| Humidity ~ | Temperature | 0.9887 |  |
| Plant cover ~ | Biomass | 0.2556 |  |
| Fungal Infection x Plant Diversity ~ | Plant Diversity | 1.47E-11 | *** |
|  | Fungi | 0.4388 |  |

**Table S7.** model fit indices of fully unconstrained model ant the final constrained model.

| model | DF | AIC | P | RMSEA | CFI | SRMR |
| --- | --- | --- | --- | --- | --- | --- |
| unconstrained | 34 | 7945.6 | 0.204 | 0.036 | 0.989 | 0.045 |
| constrained | 64 | 7912.4 | 0.361 | 0.019 | 0.994 | 0.061 |

**Table S8.**
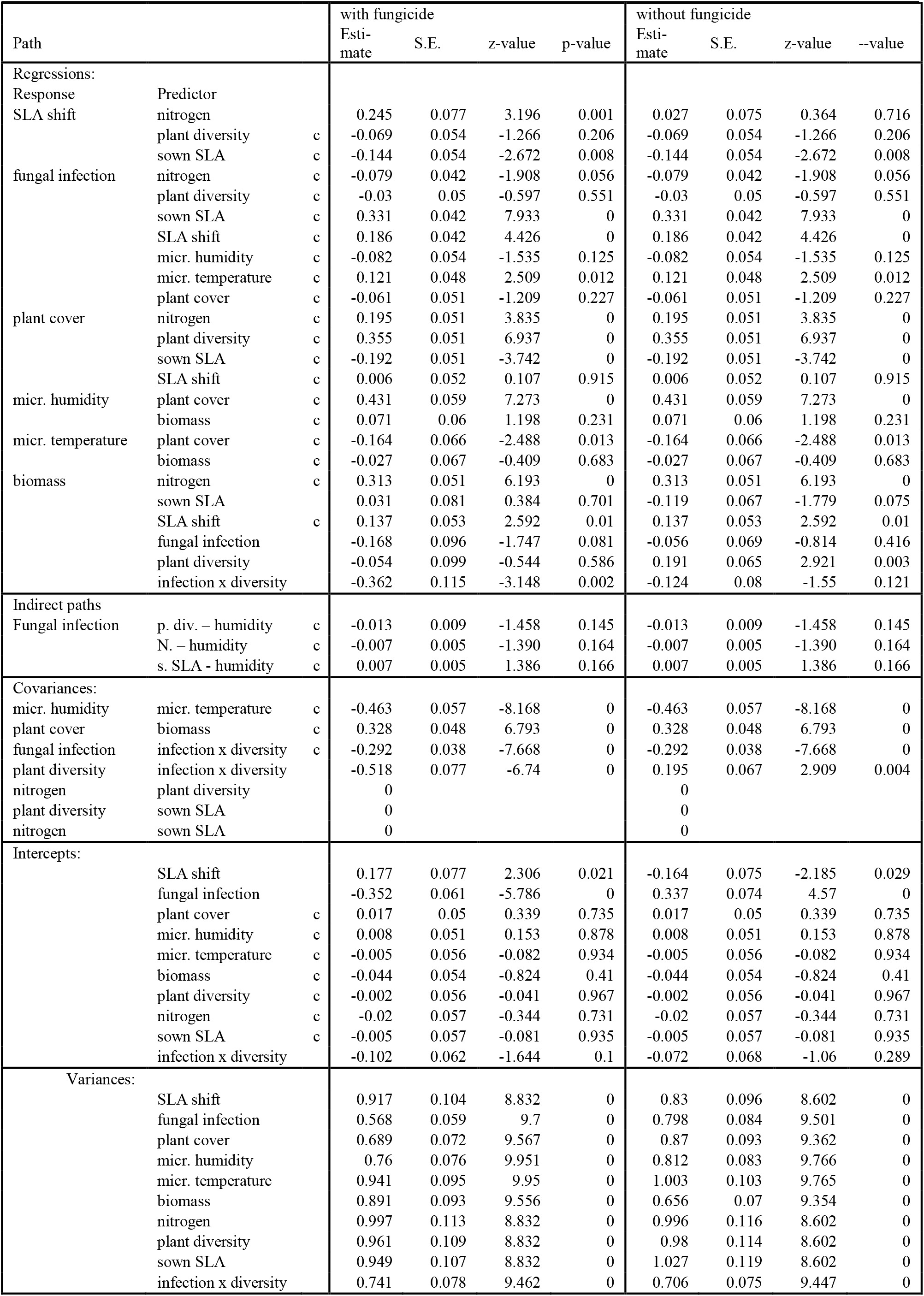
SEM path, correlation and intercept estimates with and without fungicide treatment for the standardized data. Paths/correlations/intercepts labelled with c have been constrained, because they do not significantly differ between fungicide treatments.

**Figure S6.**
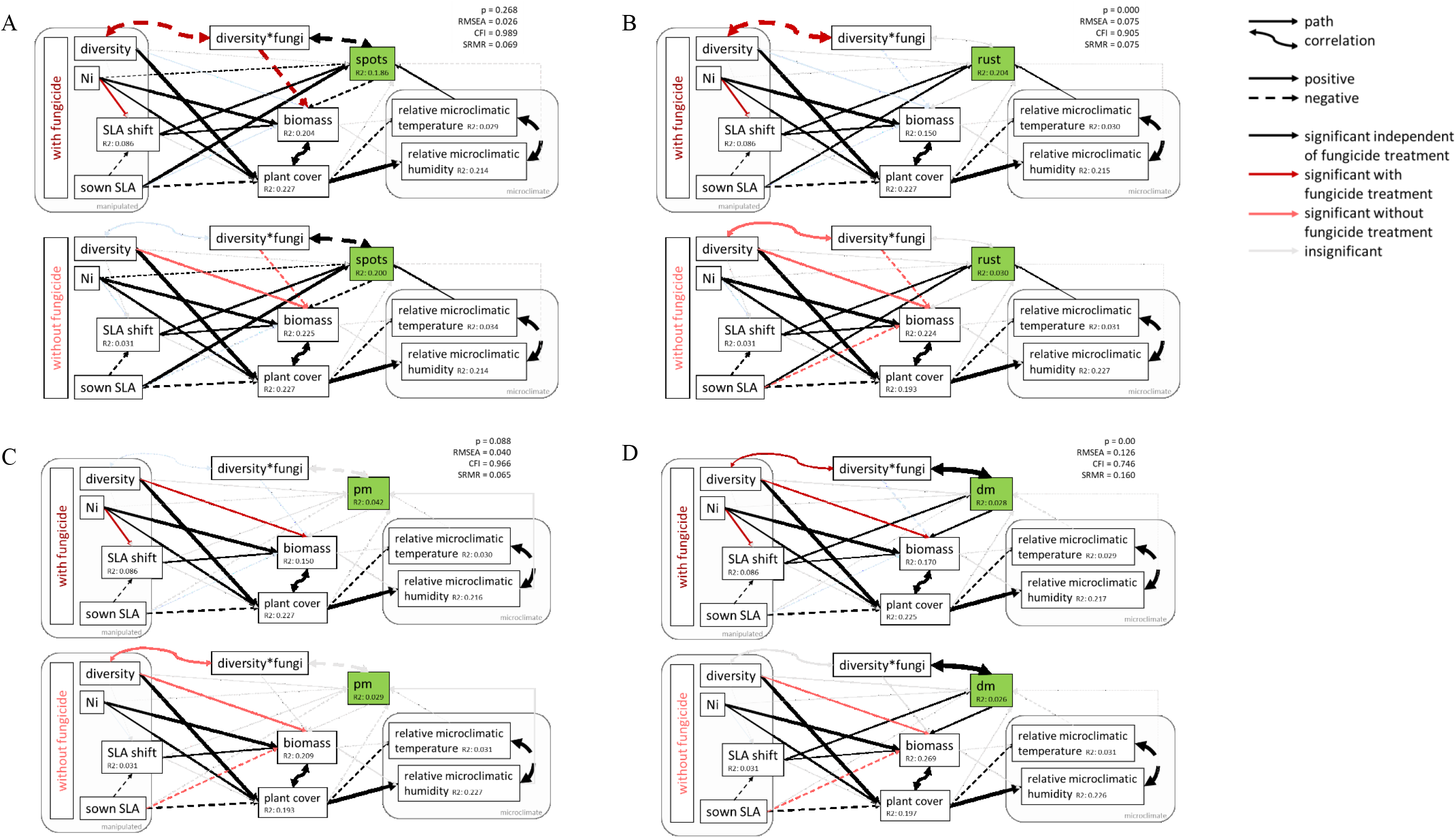
SEM results for the different fungal groups (A: leaf spots, B: rusts, C: powdery mildews, D: downy mildews)

**Table S9.** Fixed effects of the host concentration lmer (Bates *et al*. 2015), with helmert contrasts. Model: host concentration slope ∼ Nitrogen + Fungicide + Nitrogen x Fungicide + (1|Species)

| Fixed Effects | Estimate | S.E. | t-value |
| --- | --- | --- | --- |
| Intercept | 0.2262 | 0.0678 | 3.34 |
| Nitrogen | -0.0681 | 0.0505 | -1.35 |
| Fungicide | -0.1003 | 0.0505 | -1.99 |
| Nitrogen x Fungicide | -0.0006 | 0.0505 | -0.01 |
| Random Effects | Variance | SD |  |
| Species | 0.0410 | 0.2025 |  |

**Figure S7.**
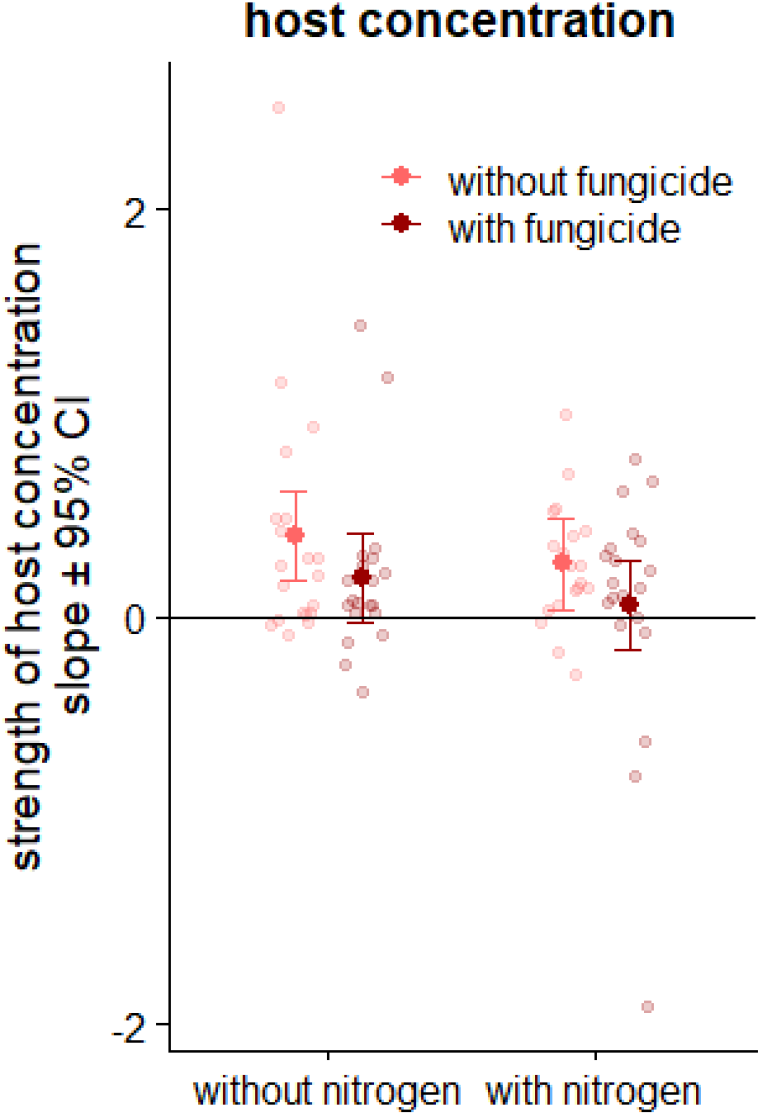
host concentration effect per nitrogen and fungicide treatment, raw data, predicted values and 95% confidence interval (obtained from the effect package in r (Fox 2003)).

